# Better late than never: the impact of hatching time on *Heterodera schachtii* parasitism

**DOI:** 10.64898/2026.05.29.728720

**Authors:** Mason Rugen-Hankey, Priya Desikan, George Harpum, Chongjing Xia, Victor Hugo Moura de Souza, Unnati Sonawala, Lida Derevnina, Beth Molloy, Anika Damm, Sebastian Eves-van den Akker

## Abstract

Plant-parasitic nematodes are a diverse, polyphyletic group of plant pathogens which can infect most plant tissues and all major crops. Amongst the most damaging clades are the cyst nematodes, which can remain dormant in the soil for decades as infection-competent, developmentally arrested, second-stage juveniles in eggs. Hatching is stimulated by a variety of factors. However, the impact of hatching factor responsiveness on nematode morphology, physiology, gene expression, and infection biology has not been explored. We examined the impact of hatching time on the beet cyst nematode, *Heterodera schachtii*. We found that late hatchers invaded host roots and established feeding sites in greater numbers than early hatchers. We demonstrate variation in baseline parasitism gene expression and in responsiveness of genes to effectostimulins, small, plant-derived molecules which upregulate parasitism genes. Three quarters of effectostimulin-induced transcriptional changes were also modulated, either positively or negatively, by hatching time. While there were no observable morphological differences between early and late hatching nematodes on the day of their emergence from the egg, the late hatchers displayed signs of faster utilisation of internal energy reserves after 7 days at 4°C, as evidenced by less body area attributed to fat, than early hatchers. Finally, we found no evidence of substantive genetic differences between early and late hatchers, they were representative of a single population, despite the observed differences in infection, gene expression, and physiology. Taken together, non-genetic differences likely drive late hatchers to more rapidly utilise their internal energy reserves, to be more responsive to host-derived signals, and to be ultimately more infective than their early hatching counterparts.

## Introduction

Plant-parasitic nematodes (PPN) are profoundly threatening agricultural pests that are estimated to cause up to 173 billion USD in crop losses annually (AbdelRazek and Balah, 2023). Like all pathogens, PPN must deliver effectors into the host to alter metabolism, immunity, and physiology to promote infection. Effectors are principally derived from two sets of pharyngeal gland cells: two subventral and one dorsal (Pellegrin and Damm et al., 2025). Some nematode species, including the major contributors to the economic losses, are sedentary endo-parasites. These are so called because they form long-term biotrophic interactions with modified cells in the vascular cylinder of the host roots (Jones et al., 2013). Over time, these nematodes may each lay hundreds of eggs which contain developmentally arrested, infection-competent, second-stage juveniles (J2s) (Siddique et al., 2022).

One of the most important groups of sedentary endo-parasitic nematodes is the cyst nematodes, which includes the genera *Heterodera, Globodera* and *Cactodera*. This group is named for its ability to contain and protect eggs within the tanned body of the deceased mother (Siddique et al., 2022). The cysts may remain dormant in the soil for many years, in ideal conditions even decades, and likely represent an adaptation to survive prolonged periods without a suitable host. Hatching of nematodes from eggs within the cyst is stimulated by a variety of factors. Root diffusates of hosts (Ochola et al., 2021), as well as a wide variety of inorganic ions including. ZnCl_2_ (Clarke and Sheperd, 1966), are typically used to induce hatching in the laboratory. J2s generally hatch after exposure to a hatching stimulant. For any given individual, exposure lasting as little as 10 minutes, or as long as several weeks, is sufficient to induce hatching. This produces a characteristic hatching curve with a ‘peak’, followed by a months-long tapered decline (Greco 1981; Zheng and Ferris 1991). Up to 90% of the total juveniles can hatch from a cyst within three weeks of exposure to hatching factors (Perry et al. 2018). Hatching curves are known to vary within populations and between species (Perry et al. 2018). It is proposed that these differences are due to hatching being fine-tuned by two mechanisms: diapause and quiescence (Perry et al. 2018). Diapause tunes hatching to cyclical, long-term conditions (such as seasons), while quiescence tunes hatching to short-term stresses (such as drought). Variation in dormancy has previously been found between cysts, across the population, throughout a growing season (Zheng and Ferris 1991). Here, we sought to examine the impact of hatching time - as opposed to time since hatching (Storey 1984) - on infection biology.

## Results

We developed a phased methodology to examine differences in hatching time, as well as time since hatching. *Heterodera schachtii* cysts (population IRS) were harvested from ~15 kg of sand and hatched in a single, large, hatching jar. Hatching was induced using 3 mM ZnCl2 solution (Clarke and Shepherd 1966), and a total of more than three million hatched J2s were collected daily for three weeks. The J2s harvested at each time-point were counted, and stored at 4 °C, and then used for a variety of experiments 7 days from when they emerged from the egg. In this way, we could determine whether time of hatching – not time since hatching – is associated with other aspects of plant-parasitic nematode infection biology.

In this experiment, the hatching curve of *H. schachtii* rapidly peaked at approximately Day 4, followed by a less rapid decline until approximately Day 9 – consistent with the literature (Greco 1981) – such that the cumulative proportion approximates a logistic growth model (Figure 1A, and Figure S1). Fifty percent of nematodes hatched within 5 days and 90% within 12 days. From this curve, two timepoints were chosen: nematodes which hatch on days 3 and 4 are referred to as the ‘early hatchers’, broadly corresponding to the peak of the hatching curve; nematodes which hatch on days 8 through 10 are referred to as the ‘late hatchers’, the final timepoint with sufficient nematodes for the analyses (184,000). The same pools of early or late hatchers were divided into three experiments (Figure 1B): experiment one measured the ability to infect white mustard (*Sinapis alba* L.); experiment two measured the transcriptional responsiveness to host-derived root extracts; and experiment three measured various morphological and physiological features.

**Figure 1.**
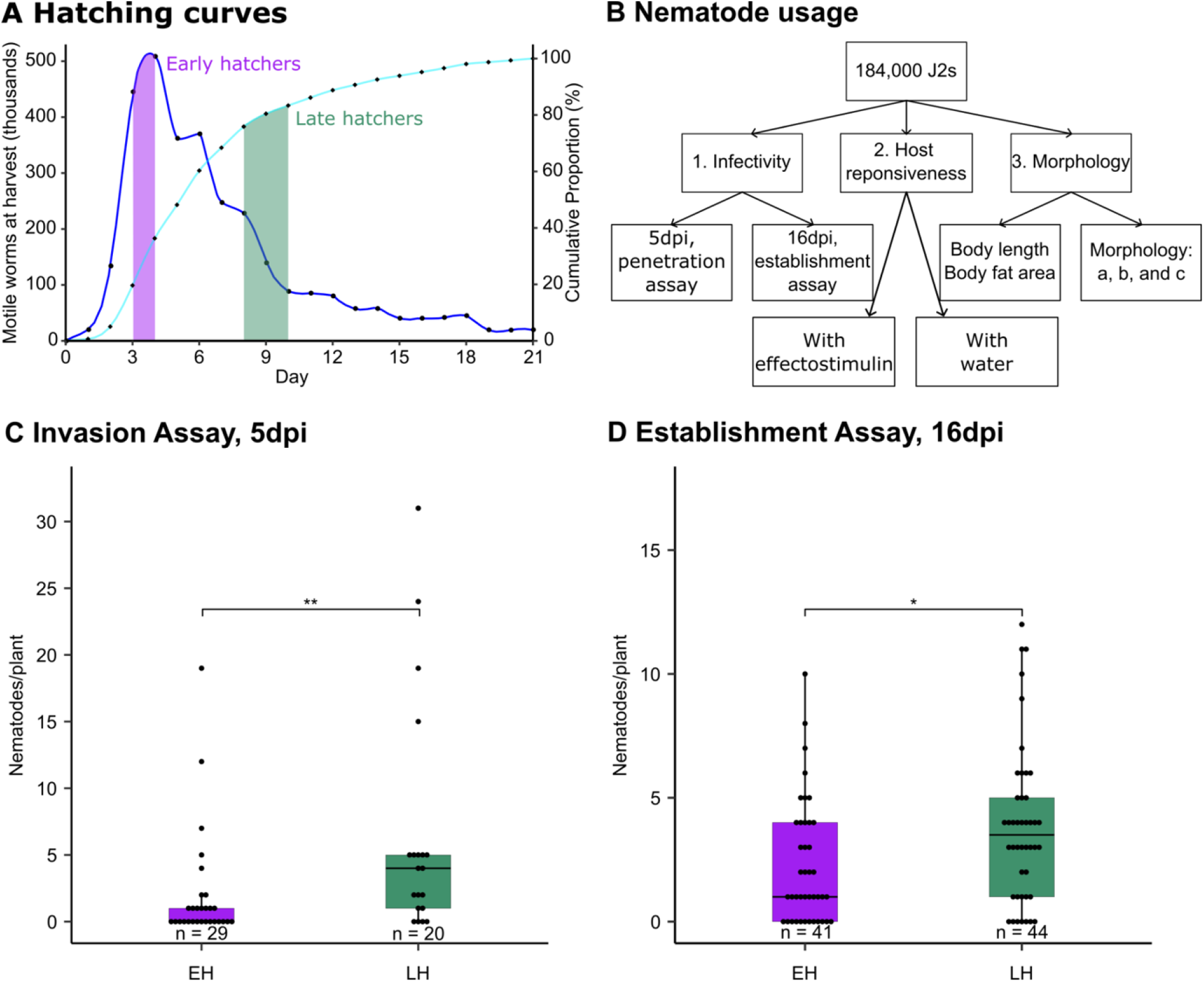
Hatching curve, downstream experiments and increased infectivity of late hatchers. **A)** Hatching curve of H. schachtii (IRS) showing number of motile nematodes hatched per day (Dark blue line, dots and left axis), and the cumulative proportion of total hatch (Light blue line, diamonds and right axis). Early hatchers (EH) denoted in purple, late hatchers (LH) denoted in green. **B)** Downstream experiments for two selected time points. Nematodes were divided into three groups and were used to assay infectivity, responsiveness to host effectostimulins, or morphological changes at harvest and after storage, across the hatching curve. **C)** Number of nematodes per plant during earlier stage parasitism – 5 dpi. **D)** Number of nematodes per plant during later stage parasitism – 16 dpi. Early hatchers shown in purple, late hatchers shown in green. Asterisks indicate statistical differences (*<0.05;**<0.01).

**Figure 2.**
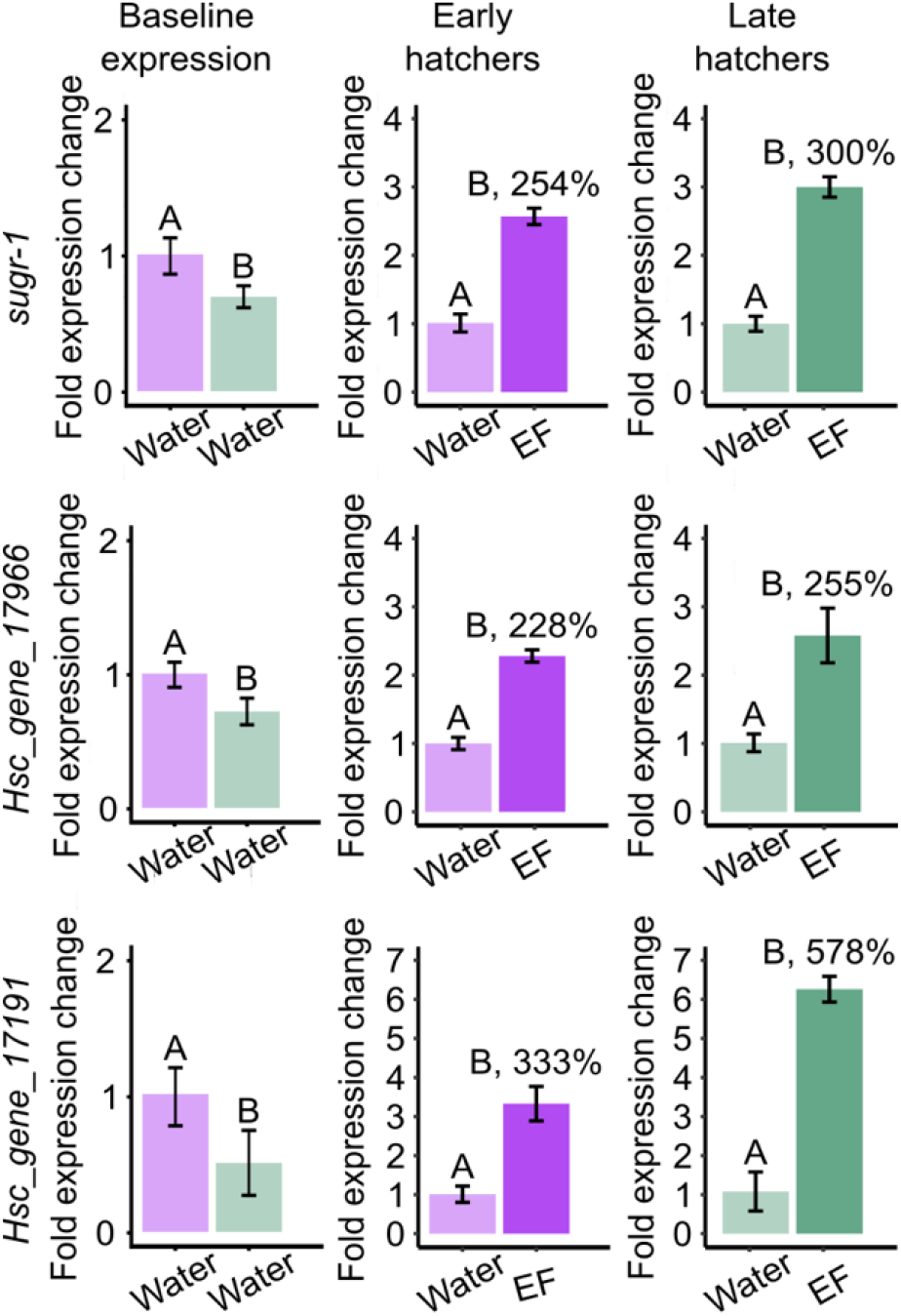
The response to effectostimulins of transcription factors associated with parasitism gene expression varies with hatching time. Left column, baseline expression between early and late hatchers. Middle column, gene fold expression changes of early hatchers, normalised to their own baseline expression for that specific gene. Right column, gene fold expression changes of late hatchers, normalised to their own baseline expression for that specific gene. Top row, sugr-1; middle-row Hsc_gene_17966; bottom-row Hsc_gene_17191. Percentages above bars indicate percentage up-regulation from the basal state. Statistical comparisons are within panels only.

To examine the impact of hatching time on infectivity, 14-day old *S. alba* were infected with 250 of either 7-day old early hatchers, or 7-day old late hatchers, at a concentration of 250 J2s / 500 μL. Forty-nine plants (n=29 and 20 for early and late hatchers, respectively) were analysed at 5 days post-infection (dpi) to measure host invasion, and the remaining eighty-five plants (n=41 and n=44 respectively) were analysed at 16 dpi to measure the establishment of biotrophy. At 5 dpi, significantly more late hatchers were observed inside host roots, compared to early hatchers (Figure 1C, Kruskal-Wallis, χ^2^ = 8.3273, df = 1, p = 0.0039). Similarly, at 16 dpi, significantly more late hatchers developed into J3 sedentary stages than early hatchers (Figure 1D, Kruskal-Wallis, χ^2^ = 5.994, df = 1, p = 0.0144). Therefore, the late hatcher advantage in invasion is maintained into early biotrophy. These data establish that late-hatching nematodes have an advantage over their earlier-hatching counterparts, particularly during host invasion.

It was recently discovered that host invasion by *H. schachtii* is controlled by the transcriptional master regulator SUGR-1, which positively regulates effectors involved in lytic functions in response to host-derived small molecules (termed effectostimulins) released from root extracts (Pellegrin and Damm et al. 2025). We therefore measured expression of *sugr-1* and two related transcription factors (TFs), Hsc_gene_17966 and Hsc_gene_17191, in early and late hatchers, in the naive state and in response to effectostimulins.

All three TFs have a lower basal expression state in late hatchers, compared to early hatchers. (*sugr1*, Welch’s t-test, t = 3.6232, df = 3.2191, p = 0.0483, FDR-adjusted; *Hsc_gene_17966*, Welch’s t-test, t = 3.7995, df = 3.9831, p = 0.0483, FDR-adjusted; *Hsc_gene_17191*, Welch’s t-test, t = 2.8024, df = 3.9473, p = 0.0494, FDR-adjusted). Late hatchers, however, were more responsive to effectostimulins and upregulated *sugr-1* more than early hatchers (300% vs 254%). The same pattern is observed for the *sugr-1* paralogue *Hsc_gene_17191* to an even greater extent (578% vs 333%). All characterised TFs are significantly upregulated after exposure to host effectostimulin (p < 0.014 in all comparisons, Welch’s t-test, FDR-adjusted).

To investigate the downstream impacts of transcription factor upregulation, we compared the global gene expression of early and late hatchers, with and without effectostimulin exposure, using whole genome RNA sequencing. Two principal components (PC1 and 2) describe 91% of the variance, as well as an apparently orthogonal relationship between variables (Figure 3A). PC1 corresponds to effectostimulin treatment and explains 72% of the variance, while PC2 corresponds to hatching time and explains 19% of the variance. Using these data, we found that 3,785 genes are differentially expressed in nematodes based on time of hatching, effectostimulin exposure, or an interaction thereof (Log2 FC>|0.5|, p<0.05).

**Figure 3.**
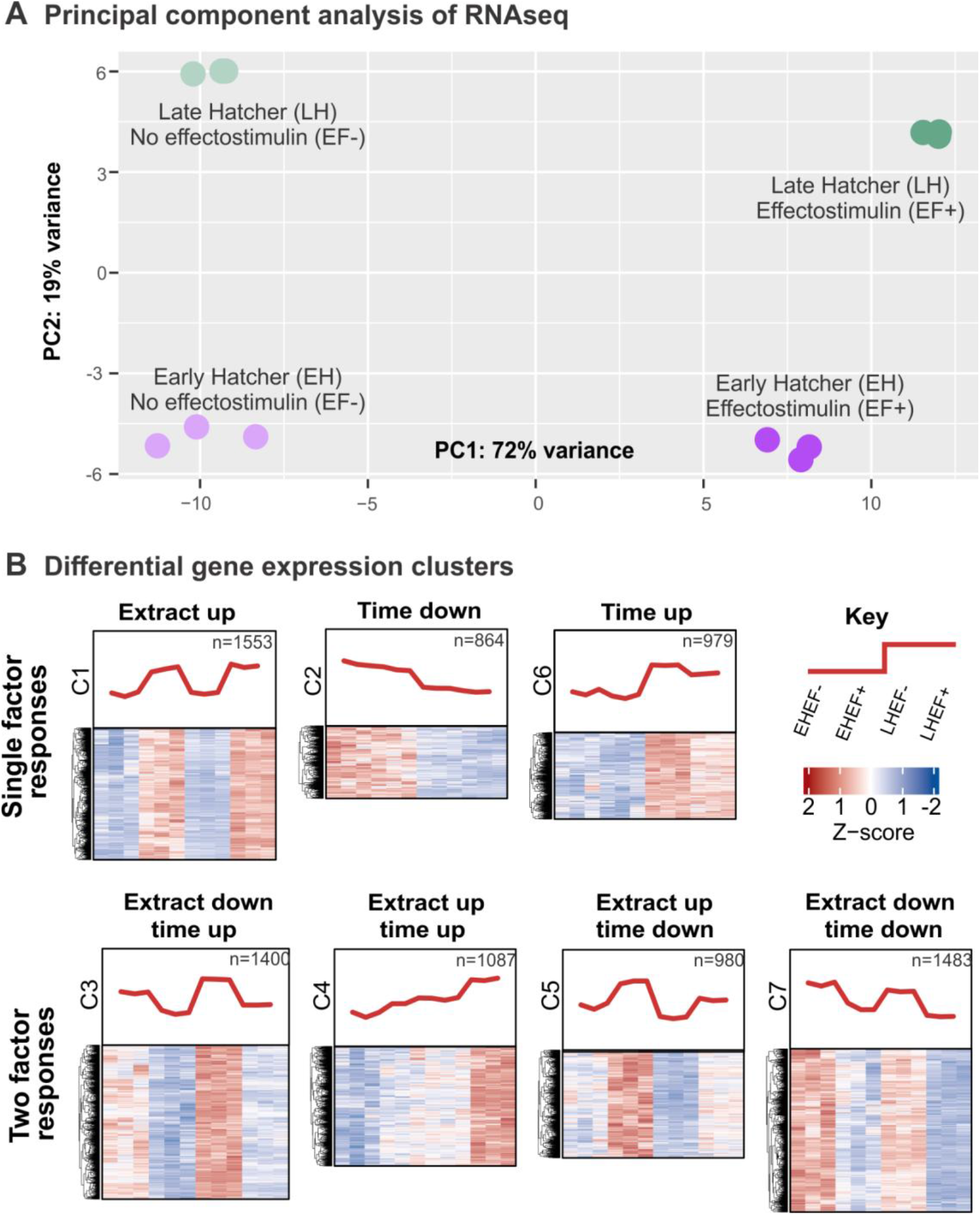
Global gene expression of early and late hatchers exposed to either effectostimulins or water. **A)** Principal component analysis of whole genome RNA sequencing following the effectostimulin or water exposure of early and late hatchers. PC1 explains 72% of the variance in gene expression and corresponds to effectostimulin exposure, while PC2 explains 19% of the variance and corresponds to the time of hatching. **B)** Clustering of differentially expressed genes according to their responses to effectostimulin exposure and time of hatching. EH EF-, early hatchers incubated in water; EH EF+ early hatchers incubated in effectostimulin; LH EF-late hatchers incubated in water; LH EF+ late hatchers incubated in effectostimulin.

All genes were clustered into seven distinctive sets (Figure 3B). Interestingly, only one cluster (cluster 1) was positively responsive to effectostimulins independent of hatching time (n=1553 genes, n=105 effectors based on the list provided in Molloy *et al*., 2025). We denoted this cluster as the “core effectostimulin response”, which represented only one quarter of all effectostimulin-responsive genes. Genes in the core effectostimulin responsive cluster 1 were enriched in effectors (hypergeometric test, p=8.74×10^−6^). However, hypergeometric tests indicate that these effectors tend not to be enriched for specific evolutionary origins, sites of production, timepoints of maximal expression and functions (Tables S1-S5).

The rest of the effectostimulin response (which is the remaining three quarters of all effectostimulin-responsive genes and two thirds of the effectostimulin activated genes) is modulated by hatching time in some way. All combinations are observed, and qualitatively assigned as Down with effectostimulins and up with hatching time (cluster 3, n=1400 genes, n=47 effectors), Up with effectostimulin and up with hatching time (cluster 4, n=1087 genes, n=126 effectors), Up with effectostimulins and down with hatching time (cluster 5, n=980 genes, n=79 effectors), and Down with effectostimulins and down with hatching time (cluster 7, n=1483 genes, n=35 effectors). According to hypergeometric tests, all except cluster 3 (Down with effectostimulins and up with hatching time) are enriched in effectors. Similarly, these effectors are not enriched for any specific properties, including evolutionary origins, sites of production, timepoints of maximal expression and functions (Tables S1-S5). Finally, changes with hatching time only are present in clusters 2 and 6, where cluster 6 is not enriched in effectors but cluster 2 is. To reconcile these observations with the principal component analysis, we surmise that while effectostimulin treatment has a larger overall effect, the effect itself is modulated in large part by hatching time. This is a remarkable contribution of an unstudied aspect of cyst nematode biology, and may indicate that various other aspects of infection biology are similarly modulated by hatching time. In summary, the expression of most effectostimulin-responsive genes is modulated by hatching time, albeit without a clear dominant regulatory pattern since all possible combinations of interactions were observed.

To understand what other aspects of parasite biology may be under control of hatching time we measured the morphology and physiology of early and late hatchers. Morphology was estimated from a random sample of at least 30 J2 nematodes for each harvest, before and after storage. For all harvests, nematode body length was measured, while the so called ‘a, b, c’s of nematode morphology (Loof 1971, a = body length / width, b = body length / oesophagus length and c = body length / tail length) were measured only for early and late hatchers. In total, more than 600 nematodes were phenotyped. Surprisingly, and somewhat peripheral to the main thrust of this study, there appears to be an increase in length after storage (Figure 4A). Lengths in aggregate were found to be significantly different between harvest and storage (Figure S2). We examined whether this trend appeared in early and later hatchers. Early hatchers increase in length after storage (Dunn’s test, p = 0.0004), as do late hatchers (Dunn’s test = 0.0313), but are not significantly different in length from one another either at harvest, or after storage (Dunn’s test, p > 0.05 in both cases). Using Principal Component Analysis, these differences can be contextualised as similar in magnitude to previously reported intraspecific differences but less than previously reported interspecific differences (Figure 4C).

**Figure 4.**
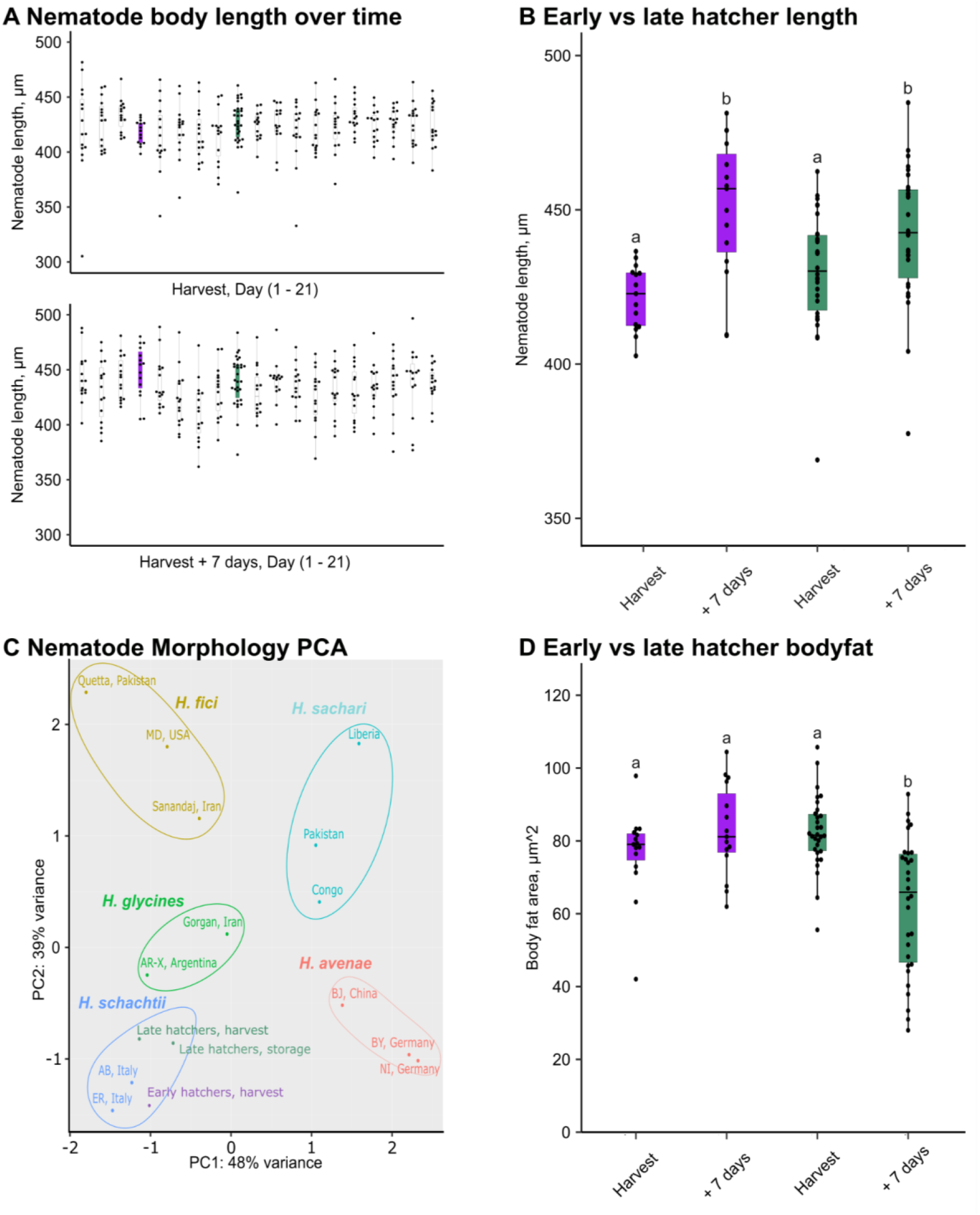
Early and late hatchers exhibit significant physiological, not morphological, differences. **A)** Lengths of randomly sampled J2s for each 24-hour period at harvest (top) and after 7-days storage (bottom). **B)** Early and late hatcher length at harvest and after 7-days storage. **C)** Principal Components 1 and 2 for nematode morphology data. Circles indicate distinct populations of nematodes or distinct timepoints. Points are coloured according to species or timepoint. Ovals are manually drawn to highlight each species grouping. **D)** Body fat area for early and late hatchers at harvest and after 7-days storage. Early hatchers in purple, late hatchers in green. Treatments with the same letter are not statistically significantly different at p < 0.05.

Most importantly, we observed characteristic signs of differential resource expenditure between early and late hatchers (Figure 4D). Early and late hatchers do not hatch with visibly significantly different lipid reserves (Figure 4D, Dunn’s test, p = 0.510). Late hatchers appear to use a significant amount of bodyfat in storage (Dunn’s test, p < 0.0001), while early hatchers appear not to (Dunn’s test, p = 0.536). Late hatchers also appear to lose fat significantly faster than early hatcher (Dunn’s test, p = 0.0019). Taken together, these data suggest that late hatchers may utilise more of their internal energy reserves than early hatchers under these conditions.

Consistent with this, cluster 3 (Down with effectostimulins and up with hatching time) and perhaps more importantly cluster 6 (Up with hatching time), are the only clusters enriched in GO terms associated with lipid metabolism/catabolism. Cluster 3: lipid catabolic process (GO:0016042) p = 0.026; lipid metabolic process (GO:0006629) p = 0.038. Cluster 6: lipid metabolic process (GO:0006629) p = 0.0066; cellular lipid metabolic process (GO:0044255) p = 0.013); lipid biosynthetic process (GO:0008610) p = 0.0160; catabolic process (GO:0009056) p = 0.0186; and organic substance catabolic process (GO:1901575) p = 0.0102.

Infectivity and gene expression differences could not be explained by morphology or physiology so we used RNA-seq reads to examine whether early and late hatchers were genetically distinct sub-populations. We selected for non-differently expressed genes (to avoid artefacts introduced by very low and very high read abundance) and calculated the Fixation Index (*F*_*ST*_) using 65,057 SNPs in cDNAs (i.e. coding and untranslated regions). A mean *F*_*ST*_ value of 1 indicates complete genetic difference, while a genetic difference of 0 indicates complete sharing of alleles between populations. We compared the *F*_*ST*_ between effectostimulin and water-treated J2s of the same timepoint (Figure 1B), which are absolutely derived from the same pool, as an internal control. We find genetic differences are extremely low between any pair of samples (*F*_*ST*_ ranges between 0.00015 and 0.0003), but actually ever so slightly higher between those we know are two halves of one and the same sample (water VS effectostimulin treatment), than those where we do not know (early vs late). We also sought to determine whether a small number of very large differences may be masked by the genome wide analyses by identifying the number of sites with *F*_*ST*_>0.25 (Figure 5B). Here we still identify no consistent differences between early and late hatchers (indeed there are still more differences between those we know are the same population). Together we conclude that there are no substantive genetic differences between early and late hatchers - they are representative of a single population, despite the observed differences in infection, gene expression, and physiology.

**Figure 5.**
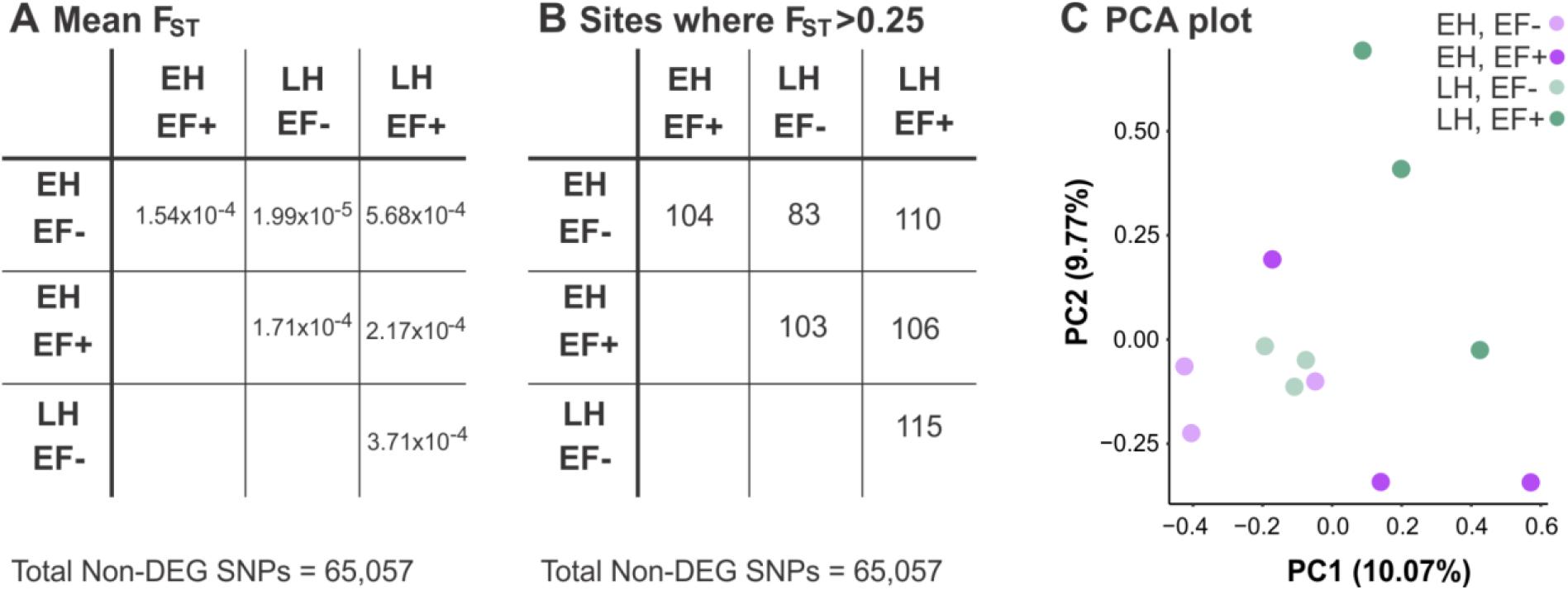
Early and late hatchers originate from a broader, genetically indistinguishable population. **A)** Mean F_ST_ across all non-differently expressed genes in early and late hatchers exposed to effectostimulin or water. **B)** Number of sites where F_ST_ exceeds 0.25 across all non-differently expressed genes in early and late hatchers exposed to effectostimulin or water. EH EF-, early hatchers incubated in water; EH EF+ early hatchers incubated in effectostimulin; LH EF-late hatchers incubated in water; LH EF+ late hatchers incubated in effectostimulin.

## Discussion

Here, we found that time of hatching significantly impacts infection biology, physiology, and host effectostimulin responsiveness of *H. schachtii*. Late hatchers invade plant roots more successfully, mobilise more of their lipid stores, and respond more to host-derived effectostimulins.

### Mechanisms for the observed differences

These individual impacts of hatching time are not necessarily causally linked to one another, only to hatching time itself. However, there are intuitive explanations for the interplay between the observations. We have correlated enriched GO terms associated with lipid metabolism in late hatchers with apparent energy reserve depletion, but have not shown causation. We hypothesise that this energy is used to boost parasitism gene expression but have not shown that this boosted gene expression increased infectivity.

Whatever the case, we find an unexpectedly large contribution of hatching time to three areas of biology: the regulation of energy reserves, the regulation of effectors, and ultimately the success of invasion. For example, while there are no gene expression clusters which are effectostimulin-responsive for late hatchers but not for early hatchers (or vice versa), cluster 4 (Up with effectostimulin and up with hatching time) is particularly interesting because it shows that the late hatchers baseline state is already at the early-hatchers stimulated state, and the later hatches stimulated state is yet higher. These data show that effector expression is regulated by multiple inputs, and more specifically that late hatchers have a mechanism to modulate host-derived signalling and certain aspects of parasitism gene expression.

### Explanations for the observed differences

If late hatchers are more infective, under what circumstances could this have been favoured by evolution? Turner and Subbotin (2013) suggested that the hatching curve is a survival strategy to avoid excessively high parasitic loads that may be fatal to both hosts and nematodes. Indeed, competition is known to reduce resource availability and increase the likelihood of developing into a male (Anjam et al 2020). As such, a broad hatching curve may strike a balance between spreading out infection while ensuring nematodes hatch when a root is present and with sufficient time in the growing season that the plant remains alive until they complete their life cycle.

In a “natural” setting, an approaching root/root system secretes hatching stimulants which are perceived by the nematodes in the eggs. It seems plausible that those nematodes which hatch early in response to hatching stimulants will have a net greater distance to travel/time to wait until they reach the root. It is intuitive to consider their slow rate of energy utilisation as adaptive in this scenario. On the contrary, the late hatchers emerge one week after in response to the same stimulus. It seems equally plausible that these nematodes have a net shorter distance to travel/time to wait as during the preceding seven days the root presumably continued its trajectory. Under these conditions, a nematode which more rapidly utilises the resources it has seems similarly adaptive.

An alternative interpretation would be one of competition between conspecifics. Early hatchers may be considered to have a head start. Late hatchers may encounter more developed plants which themselves have more resources to defend. They may have to overcome larger, tougher roots, and those with primed immunity by the interaction with their earlier hatching conspecifics. In either case, evolution may have selected late hatchers to be those that maximally invest in parasitism, more quickly utilising their resources, to “catch up” so to speak, with their early hatching conspecific. It is widely accepted that the difference in hatching behaviour of cyst and root knot nematodes represents an adaptation to their quite different host environments, host range, and survival strategies. Whether adaptive variation similarly exists between the hatching curves of different cyst nematode species, or even between populations of the same species, remains to be seen.

### The hatching curve is the sum of what whole

Hatching curves detailed within this study were characterised based on a pool of *H. schachtii* cysts extracted from approximately 15kg sand. The presence of a clear peak followed by a tapering decline can originate from one of two distinctive mechanisms. On the one hand, the hatching curve may represent an aggregated sum of similarly shaped hatching curves of individual cysts, or alternatively the sum of distinctive hatching curves of cysts within the overall population. In this second scenario, most cysts are likely to be early hatchers, while a much smaller proportion are likely to be late hatchers given the overall early skew of the hatching curve’s peak. Previous studies - albeit on distantly related cyst nematodes - perhaps support the idea of distinctive hatching profiles of individual cysts. Hatching profiles in populations of ten *Globodera pallida* or *Globodera rostochiensis* cysts exposed to potato root diffusate from the cultivar *‘*Desirée’ can sometimes be bimodally distributed depending on the temperature of hatching (Agata Kaczmarek, PhD Thesis). Hatching dynamics of individual cysts collected herein seem to suggest something in between. Regardless, genetic analyses performed here indicate that early and late hatchers do not originate from distinctive populations. Therefore, it is clear that the underlying factors shaping the hatching curve are not genetic. The gathering of these data under exposure to a single hatching stimulant indicates that this is sufficient to highlight large levels of variation in hatching dynamics across individual cysts. It is possible that simultaneous exposure to more varied hatching stimuli in the field may alter variation in the hatching profiles of individual cysts.

### Root cause of the observed differences

Regardless of whether and how the individual impacts of hatching time described here are causally linked to one another, they are causally linked to hatching time – but how? Three not mutually exclusive causes are offered: a very small number of highly impactful genetic differences, any number of epigenetic differences, and/or differences in the maternal dowry.

Even though early and late hatchers are genetically indistinguishable in this study, we cannot rule out that the responsible changes are in genes which themselves are responsive (in which case they were filtered out) but unlinked with genes harbouring the 67,057 SNPs we measured. Less unlikely, a very small number (i.e. one or two) of highly impactful genetic differences could drive the observations, masked by the overwhelming genetic similarity of the samples even when we filter for high *F*_*ST*_. In *Globodera rostochiensis*, there are three classes of calcium-binding sites on its eggshell (Perry et al. 2018). Crucially, one of these classes is thought to respond to hatching factors and induce hatching. After hatching is induced, juveniles rehydrate, become metabolically active and exit eggs. Genetic differences in eggshell-based receptors, or as yet uncharacterised downstream signalling pathways, may underlie some of the differences in hatching - but it is not obvious whether and how this difference would also explain the later observations. I.e. a very small number of very impactful genetic differences would need to explain two - as far as we are aware - unrelated phenomena: that the nematodes hatched late, and those observed differences which follow. This restriction to a small number of unlinked and impactful genetic differences, as is necessary given the absence of global differences between the populations, makes this an unfavourable hypothesis.

Alternatively, so-called epigenetic differences (e.g. histone modifications and small RNAs) would not face the same restriction. Any number could in principle be used to explain the observations. While DNA methylation is contentious in plant-parasitic nematodes, histone methylation and small RNA signalling are well established (Hassanaly-Goulamhoussen et al. 2021; Liu et al. 2023). This raises the question of when and why would these differences be deposited in this way. A first intuitive step would be to measure whether there are epigenetic differences across the hatching curve, before exploring this further.

Thirdly, there may be differences in the ‘maternal dowry’ of eggs. The egg of a nematode contains perivitelline fluid as a rich store of nutrients provided by the mother to each developing embryo (Mkandawire et al. 2022). Differences in dowry between, or within, mothers could contribute. Firstly, the hatching curve of a population is evidently the sum of many smaller curves of individual cysts. If cysts differ in dowry, that may contribute to whether it preferentially hatches early, or late. Alternatively, or in addition, given that egg production by the female begins while she is still growing, it is possible that the earliest eggs – which may preferentially correspond to early hatchers, late hatchers, or neither – receive different perivitelline fluid compositions. Given that early and late hatchers did not have observable differences in body fat composition, any differences in the maternal dowry would need to be in types, rather than amount, of nutrition. Why this would correspond to late hatching is unclear, but it is intuitive how it would correspond to a faster depletion of resources (and the observed infection and gene expression differences are intuitive to derive from that).

Taken together, we speculate that the unexpectedly large contribution of hatching time to three areas of biology - the regulation of energy reserves, the regulation of effectors, and ultimately parasitism - is likely driven by a combination of factors. It will be interesting to explore what other aspects of nematode biology are different between early and late hatchers, whether the extremes are yet more different, and whether the phenomenon is more broadly applicable to other populations, species, and pathosystems.

## Acknowledgments

Work on plant-parasitic nematodes at the University of Cambridge is supported by DEFRA licence 125034/359149/3, and funded by BBSRC grants BB/R011311/1, BB/S006397/1, BB/X006352/1, and BB/Y513246/1, a Leverhulme grant RPG-2023-001, and a UKRI FrontierResearch Grant EP/X024008/1, Royal Society research grant RGS\R1\231239, the Gatsby Charitable foundation, and through the Cambridge-Africa ALBORADA Research Fund.

## Materials and methods

### Cyst collection and hatching

*Heterodera schachtii* cysts (population IRS) were harvested from ~15 kg of infected sand using nested sieves (4000, 2000, 500, 125 and 63 µm). Cysts were then transferred onto a single layer of Miracloth (Merck), which was secured within a single large glass hatching jar (Pyrex) by a plastic ring. Cysts were covered with 75 mL of 3 mM zinc chloride solution at 21°C. Hatched nematodes were collected every 24 hours for 21 days and each harvest was stored separately in ‘Nemawash’ (0.01% v/v Tween® 20) at 4°C. Collection efficiency was improved by washing the Miracloth at each harvest using 3 mM zinc chloride in a Customised Wash Bottle (Azlon). After each collection, cysts were again covered with 75 mL of 3 mM zinc chloride solution and incubated at 21°C.

### Counting and imaging nematodes

Nematodes were collected by centrifugation (1500 x g) and the supernatant was replaced with Nemawash three times. An aliquot of nematodes was diluted in Nemawash, and the total nematode number and number of motile nematodes was counted across five or six 80 µL drops using a stereomicroscope (Leica S9D) to provide estimates for that harvest. A nematode was considered motile if it moved during the counting process. Immediately after counting, 1 mL of diluted nematode solution was added to a nematode counting slide with viewing area 18 mm x 28 mm (Astel BTU). Nematodes were bright-field imaged at 20x zoom on an inverted microscope (Oxion). Images were taken using auto-exposure on a GXCAM HiChrome-HR4 (GTVision). A minimum of 30 nematodes were imaged per time point. The remaining undiluted nematode solution was stored at 4°C for 7 days in sealed 50 mL tubes (Fisher Scientific). Nematodes were then allowed to stand at RT for 2 hours before re-counting and -imaging as described above.

### Analysing nematode morphology

Nematode morphology was measured using FIJI (v2.16.0/1.54p and build 26d66057dd) (Schindelin et al. 2012). We recorded morphology data for each harvest on the day of harvest and after 7 days storage at 4°C. Blurry nematode images were identified visually and discarded. Clear images were measured with scale 485 pixels = 100 µm. 15 nematodes per time point were analysed for: body length, body width, and body fat area (µm^2^). Body fat area and intestinal regions devoid of fat were visually identified and measured using the segmented line tool of FIJI (Figure 1C). For day 4 and days 9 and 10, a = body length / width, b = body length / oesophagus length, and c = body length / tail length – the ‘a, b, c’s of nematode morphology – were also recorded (Loof 1971).

A Principal Component Analysis (PCA) was performed using a, b, c, and nematode length. Morphology data was first normalised in R using the NormalizeTPM function of ADImpute (Leote 2025). The PCA was constructed using DESeq2 (Love et al. 2014). All external morphology data was found in Subbotin et al. (2010). The additional *Heterodera* spp. selected were chosen to represent a cross-section of the *Heterodera* genus. Species groupings were manually drawn in Inkscape.

### Treatment with effectostimulins and water

Effectostimulin extraction, J2 incubation, RNA extraction and qPCR were carried out according to the method outlined in Pellegrin et al. (2025). In brief, effectostimulin-containing root extract was prepared from *Sinapis alba* (cv. albatross). J2s were then incubated in either ultra-pure water or effectostimulins for 4 hours before flash-freezing alongside two 4 mm stainless steel bearings (ThermoFisher Scientific) in liquid nitrogen.

### RNA extraction and qRT-PCR

*Heterodera schachtii* J2s were ground to powder in a Geno/Grinder 2010 (Spex Sample Prep), and RNA was extracted according to the manufacturer’s instruction in the RNeasy Plant Mini Kit (Qiagen). cDNA was synthesised from 400 ng RNA using SuperScript IV (ThermoFisher). qRT-PCR was then performed in triplicate using LUNA Universal qPCR Master Mix (NEB) and 1 µL of cDNA. qPCR data was normalised using the Pfaffl method against two reference genes (Hsc_gene_6993 and Hsc_gene_2491) (Pfaffl 2001). Assumptions of normality and homogeneity of variances were checked visually and using Shapiro-Wilk and Levene’s tests (Levene 1960; Shapiro and Wilk 1965). All qPCR measurements were taken on distinct samples. The same RNA samples were sent for library preparation and 150 bp paired-end Illumina sequencing using the service provided by Novogene.

### Infection assay

#### Determining inoculation days

Each analysis required 184,000 J2s. The exact hatching curve for this population was unknown meaning that time points could not be pre-selected. It was therefore necessary to have plants available for each time point, and give all nematodes the same 7-day storage period. Day 4 was eventually selected to capture the variation at the peak of the curve and days 9 and 10 were selected as the latest period of the hatching curve with sufficient J2s to run the downstream experiments.

#### Growing plants

Every 24 hours, 100 *Sinapis alba* seeds (cv. albatross) in individual 15 cm seed germination pouches (mega international) were placed in one of two 24 L boxes (Really Useful Boxes) on a 16 h light/8 h dark cycle at 21 °C/20 °C in a growth chamber under white, fluorescent light (~60 µmol m^−2^ s^−1^, light level 3) (PHCBI). Plants were placed in each box on alternating days to coincide with nutrient solution change overs, reducing waste. Seed pouches were bundled into groups of 10 using 77 mm round, wavy paperclips (QConnect). Hoagland’s P10 nutrient solution (10 mM phosphate) was replenished three times per week or whenever new pouches were added to the box (Hoagland and Arnon 1950). All plants were exactly 14 days old at inoculation, with exactly 7 day old nematodes. Plants and nematodes for unused time points were disposed of.

#### Inoculation and staining

More plants were allocated to analysis at 16 days post infection (dpi), than at 5 dpi because, in our experience, nematode establishment is more variable than host invasion. A new random sample of nematodes were re-counted and re-imaged after 7-days of storage. The inoculum density was adjusted to 250 J2s/ 500 µL in Nemawash. Pouches were cut open, and the excess water was removed from the plant roots using absorbent paper. Afterwards, inoculum was carefully spread on the root system. Pouches were then closed, stacked and placed horizontally in a closed box maintained in the same growing conditions. .. After 24hours, plants were placed vertically in a box containing 2 L of Hoagland’s P10 solution, which was changed 3x weekly to minimise algal growth.

At evaluation either 5 or 16 days post-infection, plants were removed from pouches and stained using acid fuchsin according to Byrd et al (1983). Stained nematodes were counted using a stereomicroscope (Leica Stereomicroscope S9D). Number of nematodes and their relative stages were recorded. Here, nematodes recorded at 5 dpi and 16 were used to estimate invasion and establishment rates, respectively. All measurements were taken on distinct samples.

### Data analysis

#### RNA-seq data processing

Raw paired-end RNA sequencing reads were quality-filtered using Trimmomatic (version 0.39; Bolger et al., 2014) with the following parameters: LEADING:20, TRAILING:20, SLIDINGWINDOW:4:20, and MINLEN:60. This removed low-quality bases from the read ends and discarded reads shorter than 60 bp after trimming using BBduk. Trimmed reads were aligned to the reference genome of *Heterodera schachtii* (PRJNA522950.WBPS19) using STAR (version 2.7.11) in two-pass mode with parameters optimised for both gene expression quantification and variant calling. Alignments were output as coordinate-sorted BAM files with read group information included via the ‘--outSAMattrRGline’ parameter. Alignment quality metrics were assessed using mapinsights bamqc (version 1.0) and QualiMap (version 2.3)

Read counts were generated using htseq-count (version 2.0.9) at the transcript level. Expression values in Fragments Per Kilobase of transcript per Million mapped reads (FPKM) and Transcripts Per Million (TPM) were calculated using StringTie (version 3.0.0) with the -e flag to estimate abundances only for annotated transcripts.

Raw count matrices were assembled for pairwise comparisons between conditions using custom bash scripts. Differential expression analysis was performed using DESeq2 (version 1.42.1; Love et al., 2014) in R (version 4.3.2). Genes with fewer than 5 counts in at least 3 samples were filtered prior to analysis. The reference level for each comparison was explicitly set using the relevel function. Differentially expressed genes (DEGs) were identified using an adjusted p-value cutoff of 0.05 (Benjamini-Hochberg correction). Four pairwise comparisons were conducted: D4EF vs D4W, D9EF vs D9W, D9W vs D4W, and D9EF vs D4EF.

Expression clustering was performed using the ClusterGVis package (version 0.1.1) in R (version 4.5.1). TPM-normalized expression values across all 12 samples were used as input. The optimal number of clusters was determined using the getClusters function. Fuzzy c-means clustering (mfuzz method) was performed with 7 clusters over 200 iterations, applying a minimum standard deviation threshold of 0.2 to remove genes with low expression variation across samples. Genes with membership scores greater than 0.8 were retained for final cluster assignments. Cluster visualisations were generated using the visCluster function with both line plots and heatmaps.

### Genetic relationship analysis

Analysed RNA sequencing data were additionally used to investigate whether early and late hatchers can be traced to distinctive populations. For these analyses, alignments were reprocessed (adding read groups, etc) to meet GATK best practices for RNA-seq variant discovery. HaplotypeCaller function in GATK (version 4.5.0) was used to call per-sample gVCF. The 12 individual gVCF files were combined using GATK CombineGVCFs and joint genotyping was performed using GATK GenotypeGVCFs to produce a multi-sample VCF file. High-quality biallelic SNPs were filtered using VCFtools (version 0.1.18). These SNPs were further filtered in a way such that SNPs in genes that are non-differentially expression in either of the comparisons (D4EF_vs_D4W, D9EF_vs_D9W, D9W_D4W, D9EF_vs_D4EF) were selected for genetic distance and population relationship analyses. Pairwise genetic distances between samples were calculated from the filtered SNP matrix using VCF2Dis (version 1.54). Phylogenetic trees were constructed based on genetic distance matrices and visualized using standard tree visualization software.

Population structure was assessed using principal component analysis (PCA) on the filtered SNP dataset. PCA clustering was performed using VCF2PCACluster (version 1.41) with sample group information provided as input. PC1 vs PC2 scatter plots were generated to visualize genetic relationships among samples.

Population differentiation was quantified using Weir and Cockerham’s *F*_*ST*_ statistic, calculated with VCFtools --weir-fst-pop for pairwise population comparisons. SNPs with *F*_*ST*_> 0.25 were classified as highly differentiated. To test for non-random distribution of high-*F*_*ST*_ SNPs across the genome, hypergeometric tests were performed for each scaffold using a custom Python script. The hypergeometric test assessed whether the number of high-*F*_*ST*_ SNPs in each scaffold was significantly enriched compared to the genome-wide expectation.

Detailed analysis and in-house scripts are available at: https://github.com/chongjing/RNAseq_Hschachtii.

## Statistics

Statistical significance was determined using Kruskal-Wallis tests, post-hoc Dunn’s tests using Bonferroni’s Correction, and Welch’s t-tests with False-Discovery Rate corrections.

Normality was determined using Shapiro-Wilk tests. All analyses were conducted in R (version 4.2.1). All plots were generated using RStudio (v.2025.05.0+496), R (v4.2.1) and ggplot2.

For all box plots, thick black lines indicate the median; box limits indicate the 25th and 75th percentiles; whiskers extend no more than 1.5 times the interquartile range to either the largest or smallest value; data beyond the end of the whiskers are “outlying” points. Any statistical analyses were conducted separately and added via manual annotation.

## Supplemental figures

**Figure S1.**
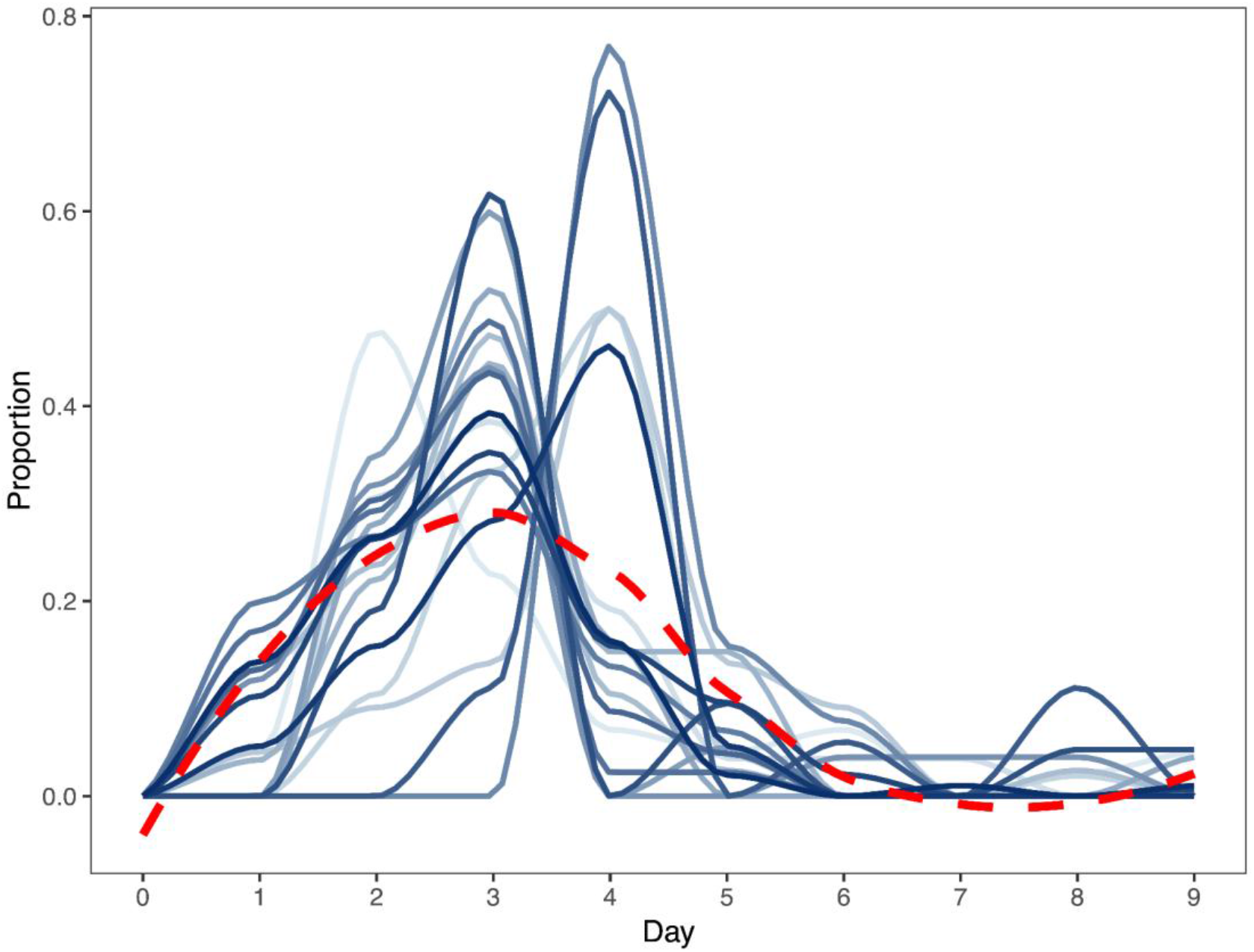
The aggregate hatching curve is seen at the single-cyst level. Individual cysts (varying blues) were harvested every 24 hours for 9 days. Proportion of cumulative hatch which hatched daily recorded. Aggregated trend (dotted red line) agrees with Figure 1A.

**Figure S2.**
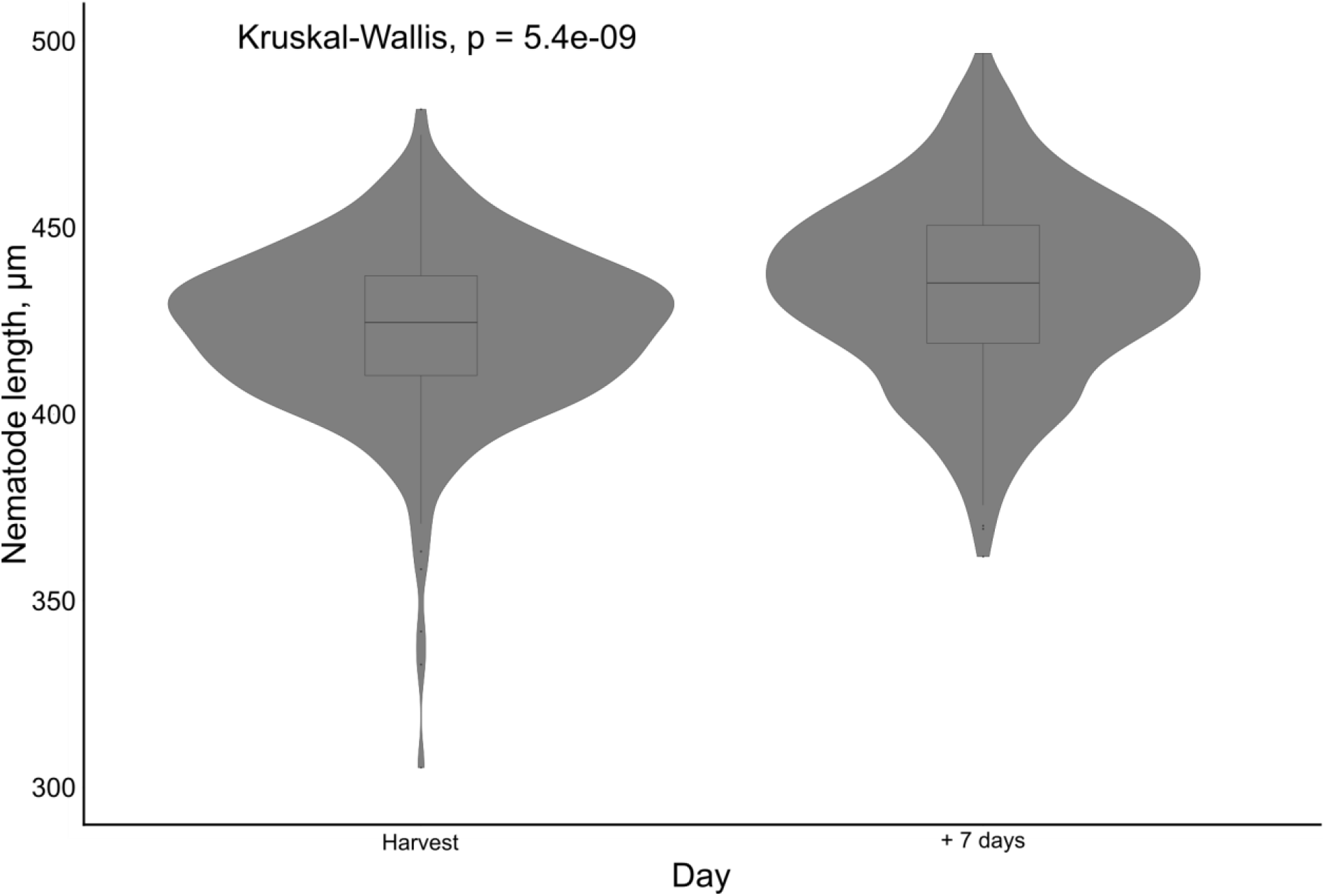
Nematode lengths at harvest and after 7 days storage, aggregated across the entire hatching curve. Nematode lengths at harvest (left) are significantly shorter than nematodes after 7 days of storage (right).

