## Supplementary material for "Better late than never: the impact of hatching time on *Heterodera schachtii* parasitism": Table S1

|  | **Cluster 1** | **Cluster 2** | **Cluster 3** | **Cluster 4** | **Cluster 5** | **Cluster 6** | **Cluster 7** |
| --- | --- | --- | --- | --- | --- | --- | --- |
| **schachtii** | 0.06 | 0.16 | 0.22 | 0.03 | 0.07 | 0.04 | 0.28 |
| **heterodera** | 0.00 | 0.10 | 0.11 | 0.06 | 0.00 | 0.09 | 0.04 |
| **cyst** | 0.05 | 0.13 | 0.17 | 0.10 | 0.12 | 0.02 | 0.19 |
| **cyst reniform** | 0.13 | 0.04 | 0.09 | 0.03 | 0.14 | 0.13 | 0.11 |
| **cyst reniform rsimi** | 0.17 | 0.20 | 0.04 | 0.01 | 0.16 | 0.18 | 0.28 |
| **cyst reniform rsimi RKN** | 0.19 | 0.13 | 0.29 | 0.12 | 0.24 | 0.01 | 0.12 |
| **down to bursaphelenchus xylophilus** | 0.17 | 0.08 | 0.29 | 0.12 | 0.27 | 0.02 | 0.38 |
| **down to ditylenchus** | 0.17 | 0.29 | 0.13 | 0.30 | 0.24 | 0.32 | 0.33 |
| **panagrolaimorpha** | 0.25 | 0.09 | 0.00 | 0.24 | 0.29 | 0.14 | 0.38 |
| **plectida** | 0.11 | 0.23 | 0.11 | 0.22 | 0.22 | 0.29 | 0.34 |
| **rhabditina** | 0.26 | 0.27 | 0.38 | 0.06 | 0.19 | 0.03 | 0.02 |
| **spirulina** | 0.40 | 0.37 | 0.66 | 0.40 | 0.37 | 0.54 | 0.03 |
| **cephlobomorpha** | 0.15 | 0.89 | 0.93 | 0.82 | 0.89 | 0.90 | 0.95 |
| **steinermatidae** | 0.39 | 0.57 | 0.25 | 0.38 | 0.35 | 0.59 | 0.20 |
| **most nematodes** | 0.21 | 0.24 | 0.08 | 0.19 | 0.24 | 0.25 | 0.03 |
| **predates nematodes** | 0.05 | 0.02 | 0.16 | 0.02 | 0.05 | 0.11 | 0.18 |

**Table S1: Probability of enrichment for effectors evolving at a specific timepoint based on hypergeometric tests.**

**Table S2: Probability of enrichment for effectors produced by a specific secretory gland based on hypergeometric tests.**

|  | **Cluster 1** | **Cluster 2** | **Cluster 3** | **Cluster 4** | **Cluster 5** | **Cluster 6** | **Cluster 7** |
| --- | --- | --- | --- | --- | --- | --- | --- |
| **SvG** | 0.18 | 0.00 | 0.04 | 0.01 | 0.16 | 0.02 | 0.31 |
| **DG** | 0.18 | 0.00 | 0.04 | 0.01 | 0.16 | 0.02 | 0.31 |

**Table S3: Probability of enrichment for effectors expressed maximally at a given timepoint based on hypergeometric tests.**

|  | **Cluster 1** | **Cluster 2** | **Cluster 3** | **Cluster 4** | **Cluster 5** | **Cluster 6** | **Cluster 7** |
| --- | --- | --- | --- | --- | --- | --- | --- |
| **10hpi** | 0.01 | 0.05 | 0.15 | 0.13 | 0.12 | 0.05 | 0.09 |
| **J2_10hpi_48hpi** | 0.10 | 0.12 | 0.19 | 0.16 | 0.21 | 0.21 | 0.01 |
| **10hpi_48hpi** | 0.08 | 0.03 | 0.15 | 0.11 | 0.09 | 0.12 | 0.16 |
| **48hpi_12dpi fem.12dpimale_24dpi** | 0.25 | 0.26 | 0.27 | 0.19 | 0.29 | 0.29 | 0.38 |
| **12dpi female_24dpi** | 0.14 | 0.10 | 0.25 | 0.18 | 0.21 | 0.29 | 0.19 |
| **48hpi** | 0.05 | 0.10 | 0.12 | 0.00 | 0.07 | 0.01 | 0.09 |
| **12dpi fem._12dpi male_24dpi** | 0.17 | 0.29 | 0.24 | 0.19 | 0.10 | 0.29 | 0.38 |
| **J2_10hpi** | 0.12 | 0.08 | 0.10 | 0.06 | 0.10 | 0.09 | 0.07 |
| **12dpi male** | 0.21 | 0.24 | 0.18 | 0.12 | 0.21 | 0.04 | 0.22 |
| **24dpi** | 0.39 | 0.57 | 0.71 | 0.38 | 0.35 | 0.59 | 0.02 |
| **12dpi fem.** | 0.29 | 0.28 | 0.19 | 0.25 | 0.28 | 0.21 | 0.46 |
| **Cyst_J2** | 0.10 | 0.01 | 0.54 | 0.34 | 0.34 | 0.39 | 0.63 |
| **10hpi_48hpi_12dpi fem.** | 0.39 | 0.34 | 0.71 | 0.41 | 0.55 | 0.33 | 0.78 |
| **12dpi fem._12dpi male** | 0.25 | 0.26 | 0.09 | 0.24 | 0.10 | 0.29 | 0.38 |
| **Not Clustered but DE** | 0.04 | 0.19 | 0.26 | 0.02 | 0.13 | 0.21 | 0.09 |
| **Decreasing** | 0.15 | 0.89 | 0.93 | 0.82 | 0.89 | 0.90 | 0.95 |
| **Cyst_J2_10hpi_48hpi** | 0.25 | 0.80 | 0.87 | 0.68 | 0.20 | 0.81 | 0.90 |
| **J2** | 0.10 | 0.39 | 0.54 | 0.29 | 0.34 | 0.17 | 0.30 |
| **Increasing** | 0.39 | 0.40 | 0.58 | 0.01 | 0.39 | 0.43 | 0.28 |
| **NA** | 0.07 | 0.29 | 0.33 | 0.17 | 0.30 | 0.33 | 0.44 |
| **J2_12dpi_male** | 0.53 | 0.31 | 0.76 | 0.40 | 0.62 | 0.29 | 0.82 |
| **J2_12dpi fem._24dpi** | 0.85 | 0.11 | 0.93 | 0.82 | 0.89 | 0.90 | 0.95 |
| **J2_48hpi_12dpi male** | 0.45 | 0.57 | 0.25 | 0.38 | 0.55 | 0.33 | 0.78 |
| **Cyst** | 0.38 | 0.51 | 0.28 | 0.22 | 0.12 | 0.54 | 0.74 |
| **10hpi_12dpi male** | 0.62 | 0.71 | 0.81 | 0.08 | 0.26 | 0.73 | 0.86 |
| **10hpi_48hpi_12dpi fem._24dpi** | 0.85 | 0.89 | 0.93 | 0.18 | 0.89 | 0.90 | 0.95 |
| **J2_10hpi_12dpi male** | 0.73 | 0.80 | 0.87 | 0.68 | 0.20 | 0.81 | 0.90 |
| **10h_48h_12d fem._12d male_24dpi** | 0.85 | 0.89 | 0.93 | 0.82 | 0.89 | 0.90 | 0.95 |

**Table S4: Probability of enrichment for effectors exhibiting cell wall degrading function based on hypergeometric tests.**

|  | **Cluster 1** | **Cluster 2** | **Cluster 3** | **Cluster 4** | **Cluster 5** | **Cluster 6** | **Cluster 7** |
| --- | --- | --- | --- | --- | --- | --- | --- |
| **Non-CWDE** | 0.15 | 0.02 | 0.01 | 0.15 | 0.11 | 0.10 | 0.29 |
| **CWDE** | 0.15 | 0.02 | 0.01 | 0.15 | 0.11 | 0.10 | 0.29 |

CWDE: cell wall degrading enzyme

**Table S5: Probability of enrichment for effectors evolving through horizontal gene transfer based on hypergeometric tests.**

|  | **Cluster 1** | **Cluster 2** | **Cluster 3** | **Cluster 4** | **Cluster 5** | **Cluster 6** | **Cluster 7** |
| --- | --- | --- | --- | --- | --- | --- | --- |
| **Non-HGT** | 0.10 | 0.03 | 0.24 | 0.18 | 0.17 | 0.06 | 0.33 |
| **HGT** | 0.10 | 0.03 | 0.24 | 0.18 | 0.17 | 0.06 | 0.33 |

HGT: horizontal gene transfer
